# Migration pulsedness alters patterns of allele fixation and local adaptation in a mainland-island model

**DOI:** 10.1101/2021.06.24.449762

**Authors:** Flora Aubree, Baptiste Lac, Ludovic Mailleret, Vincent Calcagno

**Author notes:** Author contributions: VC and FA designed the research. BL wrote a first version of the codes and made preliminary analyses. FA and VC conducted the research and wrote the manuscript with inputs from LM. Data accessibility statement: All simulation files and Java, R and Mathematica scripts are available on Zenodo (DOI 10.5281/zenodo.7133443).

## Abstract

Geneflow across populations is a critical determinant of population genetic structure, divergence and local adaptation. While evolutionary theory typically envisions geneflow as a continuous connection among populations, many processes make it fluctuating and intermittent. We analyze a mainland-island model in which migration occurs as recurrent “pulses”. We derive mathematical predictions regarding how the level of migration pulsedness affects the effective migration rate, for neutral and selected mainland alleles. We find that migration pulsedness can either decrease or increase geneflow, depending on the selection regime. Migration increases gene-flow for sufficiently (counter)selected alleles (*s* < *s*_1_), but reduces it otherwise. We provide a mathematical approximation of the threshold selection *s*_1_, which is verified in stochastic simulations. Migration pulsedness thus affects the fixation rate at different loci in opposite ways, in a way that cannot be described as a change in effective population size. We show that migration pulsedness would generally reduce the level of local adaptation, and introduce an additional genetic load: the “pulsedness load”. Our results indicate that migration pulsedness can be detrimental to the adaptation and persistence of small peripheral populations, with implications in management and conservation. Our results highlight temporally variable migration as an important process for evolutionary and population genetics.

## Introduction

Geneflow between populations, as a major determinant of evolutionary dynamics, has received considerable interest since almost a century. Depending on its intensity and interactions with other evolutionary forces, geneflow has a range of contrasting effects (e.g. Felsenstein, 1976; Lenormand, 2002; Bürger, 2014; Tigano & Friesen, 2016). In a focal population, it can enhance genetic diversity, prevent inbreeding, or on the contrary hamper local adaptation (Gomulkiewicz *et al.*, 1999; Garant *et al.*, 2007; Bürger & Akerman, 2011). Across populations, it controls the spatial spread of novel mutations, the maintenance of polymorphisms, the level of population divergence and, eventually, the possibility of speciation (e.g. Maynard-Smith, 1966; Johnson *et al.*, 2000; Yeaman & Otto, 2011; Mailund *et al.*, 2012; Rousset, 2013; Feder *et al.*, 2019).

The consequences of geneflow have been studied in a range of spatial configurations, such as two interconnected populations (Maynard-Smith, 1966; Yeaman & Otto, 2011), mainland-island systems (Johnson *et al.*, 2000; Bürger & Akerman, 2011) or metapopulations (Slatkin, 1981; Rousset, 2013; Feder *et al.*, 2019). However, temporal variability in the flows of propagules is rarely investigated in theoretical studies, and those usually consider the process of migration (here used in the sense of dispersal) as constant. This is unfortunate, as migration is often governed by time-fluctuating and potentially highly unsteady phenomena that would make it temporally variable (Peniston *et al.*, 2019).

Causes of temporal variability in migration rate are almost endless. Among the most frequently cited phenomena, a first category relates to environmental variations. Geographical barriers can change depending on land bridges (Morris-Pocock *et al.*, 2016; Keyse *et al.*, 2018), sea levels fluctuations (Hewitt, 2000) or habitat fragmentation (Peacock & Smith, 1997). Dispersal can also be affected by variation in oceanic and atmospheric currents (Renner, 2004; White *et al.*, 2010; Smith *et al.*, 2018; Benestan *et al.*, 2021; see also Catalano *et al.*, 2020 for a recent attempt to quantify dispersal variability). Last, but not least, dispersal is also affected by extreme meteorological or climatic events such as floods and storms (Reed *et al.*, 1988; Boedeltje *et al.*, 2004) that can cause rafting events (Garden *et al.*, 2014; Carlton *et al.*, 2017). These extreme phenomena are bound to become more prevalent with climate change (Masson-Delmotte *et al.*, 2018). A second category includes variations caused by species life-history traits, such as mast seeding, ballooning (Bishop, 1990) or various forms of group or clump dispersal (Soubeyrand *et al.*, 2015). And finally, a third category encompasses variation related to dispersal by animal vectors (Yamazaki *et al.*, 2016; Martin & Turner, 2018) or human activities (through ballast waters for instance, see Carlton & Cohen, 2003). All three categories are known to result in variable dispersal rates, in the form of random fluctuations (Yamazaki *et al.*, 2016), intermittent flows (Hewitt, 2000; Keyse *et al.*, 2018) or episodic bursts of migration (Peacock & Smith, 1997; Reed *et al.*, 1988; Carlton & Cohen, 2003; Reiners & Driese, 2004; Morris-Pocock *et al.*, 2016). The latter form is particularly common and is often referred to as “pulsed migration” (Boedeltje *et al.*, 2004; Bobadilla & Santelices, 2005; Folinsbee & Brooks, 2007; Smith *et al.*, 2018; Martin & Turner, 2018).

The relatively few existing theoretical studies suggest that migration variability can have important population genetics and adaptive implications. Nagylaki (1979), Latter & Sved (1981) and Whitlock (1992) have all used discrete-time island models and considered neutral genetic variation only (infinite alleles model). All three concur in predicting that temporal variability in migration rates should decrease effective geneflow, and thus increase differentiation among populations (see also Rousset, 2013). In a two-population model, Yamaguchi & Iwasa (2013), and following papers, studied the fixation of incompatible mutations and the progress to allopatric speciation, and found that speciation occurred faster with variable migration. Some studies have investigated non-neutral cases in spatially heterogeneous environments, in particular in the context of source-sink population systems. Gaggiotti & Smouse (1996) found that spacing out migration events, while keeping the mean number of migrants constant, decreases the level of genetic variation in the sink population. Rice & Papadopoulos (2009) studied a mainland-island model and suggested that neglecting migration stochasticity would generally lead to overestimating the impacts of migration on adaptation. More recently, Peniston *et al.* (2019) extended the results of Gaggiotti & Smouse (1996), investigating the impact of temporally pulsed migration on the level of local adaptation in the sink population. They found that spacing out migration events (with a constant mean number of migrants) can either hamper or facilitate adaptation in a harsh sink environment, depending on genetic scenarios. In a different context, Matias *et al.* (2013) studied specific diversity in a metacommunity model, and found that with randomly fluctuating migration rates, larger mean dispersal values were needed to produce the same local species richness. Overall, these studies seemingly converge on a negative impact of dispersal variability on effective migration rate. However they remain few and limited in their scope. Most of them underline the need for more attention being given to the consequences of migration variability (Peniston *et al.*, 2019).

The present study aims at providing a more comprehensive appraisal of the consequences of temporal variability in migration, considering both neutral, beneficial and deleterious alleles, under various forms of selection and levels of dominance. We will consider a mainland-island system where a small island population receives migrants from a large mainland population (Wright, 1931; Felsenstein, 1976; Bürger, 2014). In the island population, we will model the dynamics of allele frequencies in continuous time, at one locus, in diploid individuals. We will focus on the case of “pulsed” migration patterns, i.e. geneflow from the mainland occurs as bursts of migration of variable size and frequency (e.g. Rice & Papadopoulos, 2009; Peniston *et al.*, 2019). We explore the range of migration “pulsedness” from continuous migration (independent migration of individuals) to very pulsed migration (rare migration of groups of individuals).

Mathematical predictions for how the level of migration pulsedness should affect the effective migration rate are derived under a low-migration limit. These predictions are validated in stochastic simulations with large migration rates and logistic population dynamics on the island. We show that for neutral alleles, migration pulsedness reduces the effective migration rate, thus decreasing the rate of allele fixation, which is an accordance with earlier theoretical results. However, under general scenarios of selection, this conclusion can change quantitatively, and even reverse. We find that if the selection coefficient falls below some negative threshold value *s*_1_, of which we provide an analytical expression, the qualitative impact of migration pulsedness switches to positive, so that the effective migration rate increases for those deleterious alleles. Moreover, for sufficiently recessive deleterious alleles, migration pulsedness may have a non monotonous effect on the effective migration rate. The interplay of selection, dominance and drift is thus found to play an essential role in determining the impact of migration variability on effective migration rate.

Our results show that the effect of migration pulsedness is not uniform across loci subject to different selection regimes and dominance levels. Over an entire genome, migration pulsedness effectively homogenizes the fixation rates across loci, in a way that cannot be simply described as a change in effective population size. This has consequences for the dynamics of mean fitness and local adaptation, by creating an additional migration load that we call the “pulsedness load”. Overall, our results highlight migration variability as an important property to be taken into account when studying geneflow and local adaptation. Specifically, pulsed migration patterns could overall be detrimental to the local adaptation and persistence of small peripheral populations.

## Methods

### Mainland-island model

A large mainland population is connected to a small island population of finite size *N*, and migrant individuals flow from the former into the latter. We consider arbitrary loci at which some allele *A* is fixed in the mainland whereas a different allele *a* was initially fixed in the island population (fig. 1). Individuals are diploid (there are 2*N* genes at any locus) and mate randomly within the island population. The three diploid genotypes *AA*, *Aa* and *aa* have fitness values 1 + *s*, 1 + *hs* and 1, with *h* the degree of dominance and *s* the selection coefficient, as usually defined (Thurman & Barrett, 2016). Results directly extend to haploids if one considers genic selection (*h* = 0.5) and takes the number of genes to be *N*. For simplicity, mutation is neglected. The evolution of allele frequencies in the island population is modelled in continuous time (overlapping generations, Moran, 1962).

**Fig. 1.**
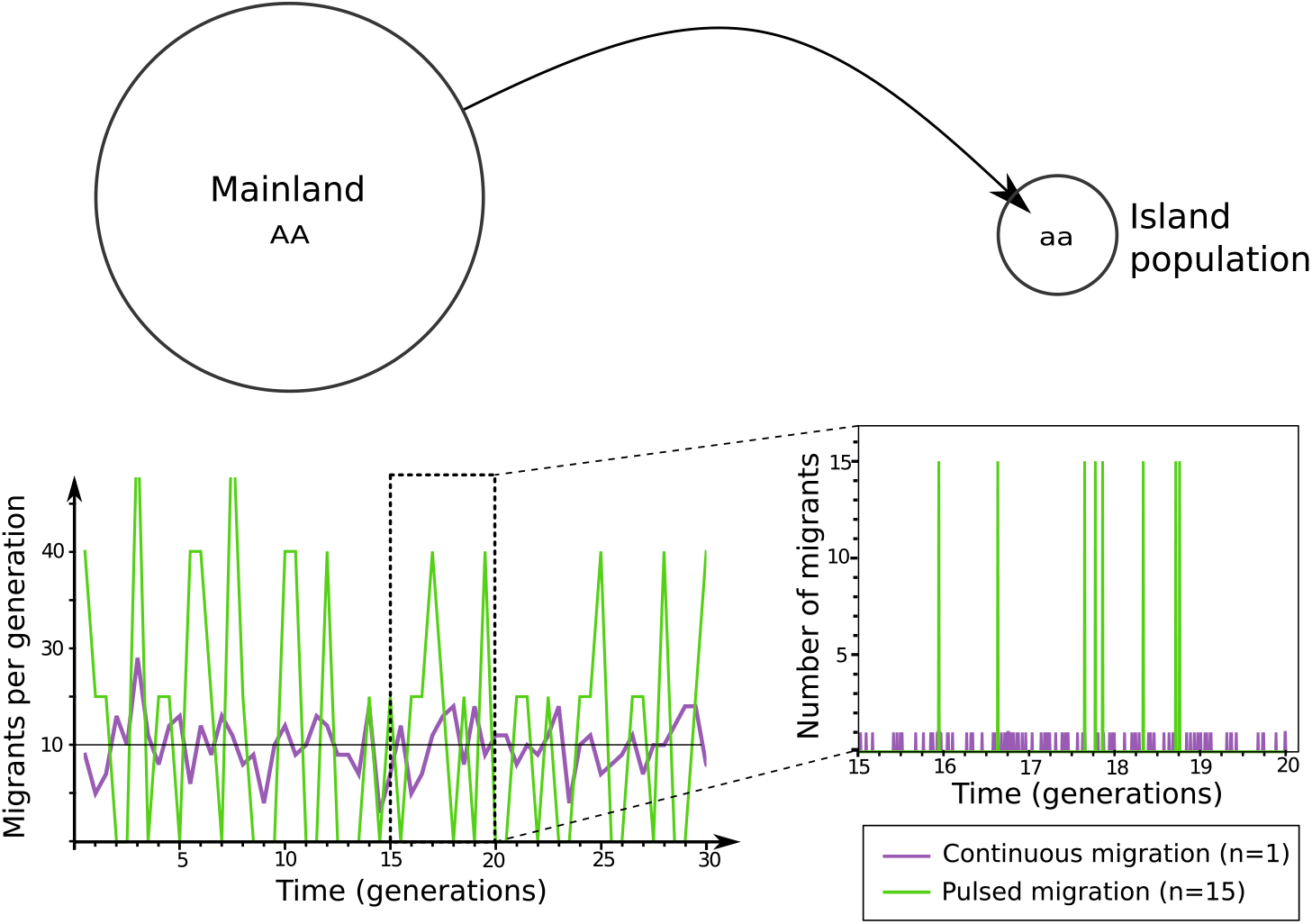
Number of migrants as a function of time in the continuous case (*n* = 1, independent migration of individuals) and in a pulsed migration case (*n* = 15). The left panel averages the number of migrants over one generation. The right panel is a close-up over 5 generations, showing the arrival of individual migrants at each migration event. Illustrative parameters: *m* =10, *T* = 1.

### Modelling of pulsed migration

Migration occurs as discrete migration events (pulses of migrants) corresponding to the arrival of *n* individuals into the island population (see Yamaguchi & Iwasa, 2013, for a similar description of migration). The overall intensity of migration is controlled by the migration rate *m*, the mean number of migrants per unit of time. To keep this mean number of migrants constant, we impose that the more individuals arrive per migration event (the larger *n*), the less frequent migration events are (Peniston *et al.*, 2019). Specifically, the rate at which migration events occur is taken to be *m/n*. If *n* =1, individuals migrate independently, corresponding to the classical continuous form of migration. When *n* exceeds one, migrants arrive more sporadically, but in larger packs. We will thus increase *n* to make migration more and more pulsed, and evaluate how this affects predictions compared to continuous migration (*n* = 1).

Parameter *n* controls the degree of migration pulsedness. More precisely, the number of migration events that occur in one generation time (*T*) follows a Poisson distribution with mean *mT/n.* Therefore *mT/n* migration events occur on average, with a variance equal to *mT/n*. It follows that the variance in the number of migrants (*n* × *mT/n*) per generation is *n*^2^*mT/n* = *nmT*. Parameter *n* thus quantifies the degree of overdispersion (variance inflation) in the number of migrants per generation, relative to the continuous case. Migration events can range from very frequent and of small intensity, with minimal temporal variance in the number of migrants per generation (low *n*), to more infrequent and intense, with large temporal variance (large *n*; Fig. 1).

### Metrics of interest

In such a mainland-island model, fixation of mainland alleles in the island eventually occurs with probability one, but we are interested in how long fixation takes, depending on the pulsedness level *n*. In other words, we seek to understand whether migration pulsedness, all else equal, is favorable or unfavorable to geneflow. We want to know this for mainland alleles with different selective values (*h* and *s*), and across different population sizes (*N*). To compare pulsed migration with continuous migration, we will compute the effective migration rate *m_e_*, defined as the migration rate that would be required to produce the same time to allele fixation, all else equal, under continuous migration (i.e. with *n* = 1). This definition is a variant of others (Wang & Whitlock, 2003; Kobayashi *et al.*, 2008; Rice & Papadopoulos, 2009), adapted to the study of migration pulsedness. The value of *m_e_* is by definition *m* when *n* = 1, and can deviate from *m* for larger *n*: if *m_e_* > *m*, migration pulsedness promotes geneflow, whereas if *m_e_* < *m*, it reduces geneflow. Following Kobayashi *et al.* (2008), we will further compute the geneflow factor, *m_e/m_*: if the geneflow factor is greater than one, migration mulsedness promotes geneflow, whereas if it is less than one, it reduces geneflow.

Note that all comparisons are done for a given locus with a given selective regime, not across different selective regimes. Obviously, beneficial alleles go to fixation faster than neutral or maladaptive ones, for any kind of migration. Hence our effective migration rate and geneflow factor are not “relative to a neutral allele”, as sometimes used, but rather “relative to continuous migration, for an identical allele” (Kobayashi *et al.*, 2008).

### Stochastic simulations

We simulated the island population as a continuous time birth-death-immigration process, using a Gillespie algorithm. The population dynamics of the island population followed a stochastic logistic model with carrying capacity *K* (Goel & RichterDyn, 1974). Each diploid individual has some basal death rate *d*, birth rate *b* = *d*, and density dependence acts through the death rate: total death rate is 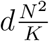 while total birth rate is *bN*. Selection occurs at reproduction (fertility selection): reproducing individuals are picked at random with odds proportional to their selective values. For computational efficiency, we used an optimized Gillespie algorithm (see Supplementary Information; Section 1 for a full description of the simulation algorithm).

The island is initially fixed for allele *a* and at carrying capacity (*N* = *K*). At the beginning of a simulation, it starts receiving *AA* migrants from the mainland, under the stochastic migration process described above (Fig. 1). The simulation ends when allele *A* gets fixed in the island. We conducted 10,000 replicates per parameter set to capture the stochasticity in migration times, population size fluctuations, and genetic drift. Effective migration rates were determined from the observed time it takes for the mainland allele to get fixed in the island, using a pre-computed abacus (S.I.; Section 1.3).

### Mathematical approximation

To allow analytical progress, we make the standard assumption that population regulation is fast, so that population size is effectively constant at *N* in the island. However, we cannot employ the equally standard method of diffusion approximations. Indeed, the latter would require *n* to be smallish relative to *N*, *de facto* preventing the description of pulsed migration patterns (see Kimura, 1962; Yamaguchi & Iwasa, 2013). We rather use an alternative approximation, assuming that migration events are rare enough, so that genetic drift and selection result in allele fixation before the next migration event occurs.

With these assumptions, at each migration event, *n* homozygous *AA* migrants arrive into the island population containing either *N aa* individuals (if fixation did not occur yet) or *N AA* individuals (if it did). Following every migration event, population regulation quickly takes population size back to *N*. Until fixation of the mainland allele, each migration event brings the latter in initial frequency 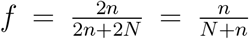, and the outcome of the particular migration event is either fixation (success), or disappearance (failure) (see also Yamaguchi & Iwasa, 2013). The probability of success can be well approximated by the fixation probability of an allele in initial frequency *f* in a closed population of size *N_e_, u* (*f, N_e_, s,h*). Expressions of *u* are are well-known (Kimura, 1962; Whitlock, 2003). For simplicity, we will typically consider *N_e_* is equal to *N* in computations, even though *N_e_* could often be smaller than population size, including in our simulations (owing to demographic fluctuations). To simplify notations, we’ll write *u* (*f*) or *u* (*f,N_e_, s, h*).

From this, we can derive the probability that the mainland allele has not yet swamped the island after a given time, and the mean time before it does, from an exponential distribution with rate –*u* (*f*) *m/n* (S.I.; section 2.2).

### Parameter values

The island population should be sufficiently isolated, otherwise it will immediately be swamped by the mainland and geneflow is a trivial issue. A famous rule-of-thumb is the so-called “one migrant per generation rule” (Mills & Allendorf, 1996, see also Blanquart *et al.*, 2013), stating that migration is strong if more than one migrant enters the population every generation. Without loss of generality, we’ll set the basal death rate as *d* = *N* in our simulations, so that the generation time (defined as the average time for *N* deaths to occur, see Moran, 1962) is *T* ≈ 1. Since the mean number of migrants per generation is *mT*, this means that, roughly speaking, *m* should not exceed 1. In turn, our mathematical approximation requires the average time between migration events (*n/m*) to be much larger than the average time to fixation after one event. Using classical theory (Kimura & Ohta, 1969; Whitlock, 2003; Otto & Whitlock, 2013), we can show that this always holds if 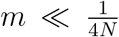. This is a sufficient condition: the approximation should hold well even for larger migration rates (S.I.; section 2.1).

We will consider four scenarios regarding the intensity of migration in simulations: (i) *very low* migration (*m* = 0.001); (ii) *low* migration (*m* = 0.1); (iii) *medium* migration (*m* =1), and (iv) *strong* migration (*m* =10). The first scenario will be used to check that simulations do converge to our mathematical approximation. The three other scenarios span the one migrant per generation rule by one order of magnitude in both directions. The last scenario (strong migration) is the one illustrated in fig. 1.

The level of migration pulsedness *n* will be systematically varied between 1 and *N*, the latter being an extreme case such that migration pulses temporarily double the local population size and take migrants at 50% frequency. Selection parameters will also be systematically varied in plausible ranges: *s* between –0.1 and 0.1 (Martin & Lenormand, 2015; Thurman & Barrett, 2016), and *h* between 0 and 1. The population size *N* will be varied between 50 and 200 (Palstra & Ruzzante, 2008; Peniston *et al.*, 2019). With such parameters, the average proportion of immigrant individuals per generation (*m*/(*N* + *m*)) varies between 0.0005 and 0.2.

## Results

### Mathematical predictions

#### General criterion for the impact of migration pulsedness

Under our mathematical approximation, the rate of fixation of a mainland allele is the product of *m/n* and the probability of success 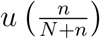 of a particular migration event (see above), and the mean time to fixation is the inverse of this rate. By definition of the effective migration rate, we must have 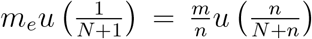. It follows that in order to have a geneflow factor greater than one (*m_e_* > *m*, i.e. migration pulsedness increases geneflow), we must have:

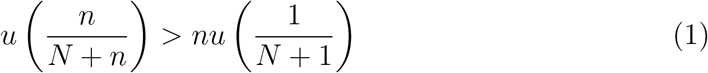

This condition yields a simple graphical criterion to determine whether migration pulsedness promotes or decreases geneflow, based on the shape of the *u* function corresponding to any particular selective regime. It also provides an operational way to derive quantitative mathematical predictions for specific types of genetic variation, as we’ll proceed to do now.

#### Neutral variation

At a locus where the mainland allele is neutral, we can apply criterion (1) using the fixation probability of a neutral allele, which is well known to be its initial frequency:

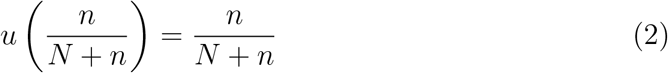

In that case, it is straightforward to see that criterion (1) is never met, for any *n*. Note that in the above equation, *N* is the actual census population size, not *N_e_*, even if they differ. We thus predict that migration pulsedness should invariably decrease the effective migration rate, i.e. decrease geneflow. The more pulsed migration is, the larger the effect is.

#### Selected variation

In the general case of selection with dominance, an expression for function *u* is:

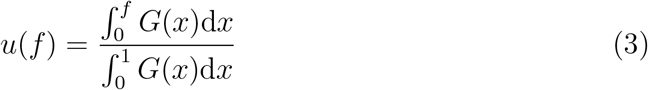

with *G*(*x*) = exp [2*N_e_s*((2*h* – 1)*x*^2^ – 2*hx*)] (Kimura, 1962).

Note that in the expression of *G*(*x*), the effective population size *N_e_* matters, if different from the census population size *N*.

This expression allows much greater flexibility in the shape of the function. Function *u*(*f*) is well known to be either strictly concave, strictly convex, or convexconcave with an inflexion point, depending on *s* and *h*. A particular case is the additive case (frequency-independent selection in haploids, or codominance in diploids, i.e. *h* = 1/2), for which a simpler explicit solution exists (Kimura, 1962; Maruyama, 1970; Whitlock, 2003).

Mathematical predictions based on numerical analyses for all *s* and *h* values are presented in fig. 2. Even though eq. (3) is much less mathematically tractable, we can still make some analytical progress. Criterion (1) can be viewed as a comparison to linearity for function *u*[*f*(*n*)] (S.I.; section 2.3): if initially (for low levels of pulsedness, i.e. *n* close to 1), the function is concave, then the criterion will not be met. If on the contrary it is convex, then the criterion will be met. We can therefore determine whether making migration pulsed increases or decreases geneflow by studying the curvature of *u*[*f*(*n*)] at *n* =1. Since *f* is a concave function of *n*, we know that *u*[*f*(*n*)] will be more prone to being concave than *u*(*f*).

**Fig. 2.**
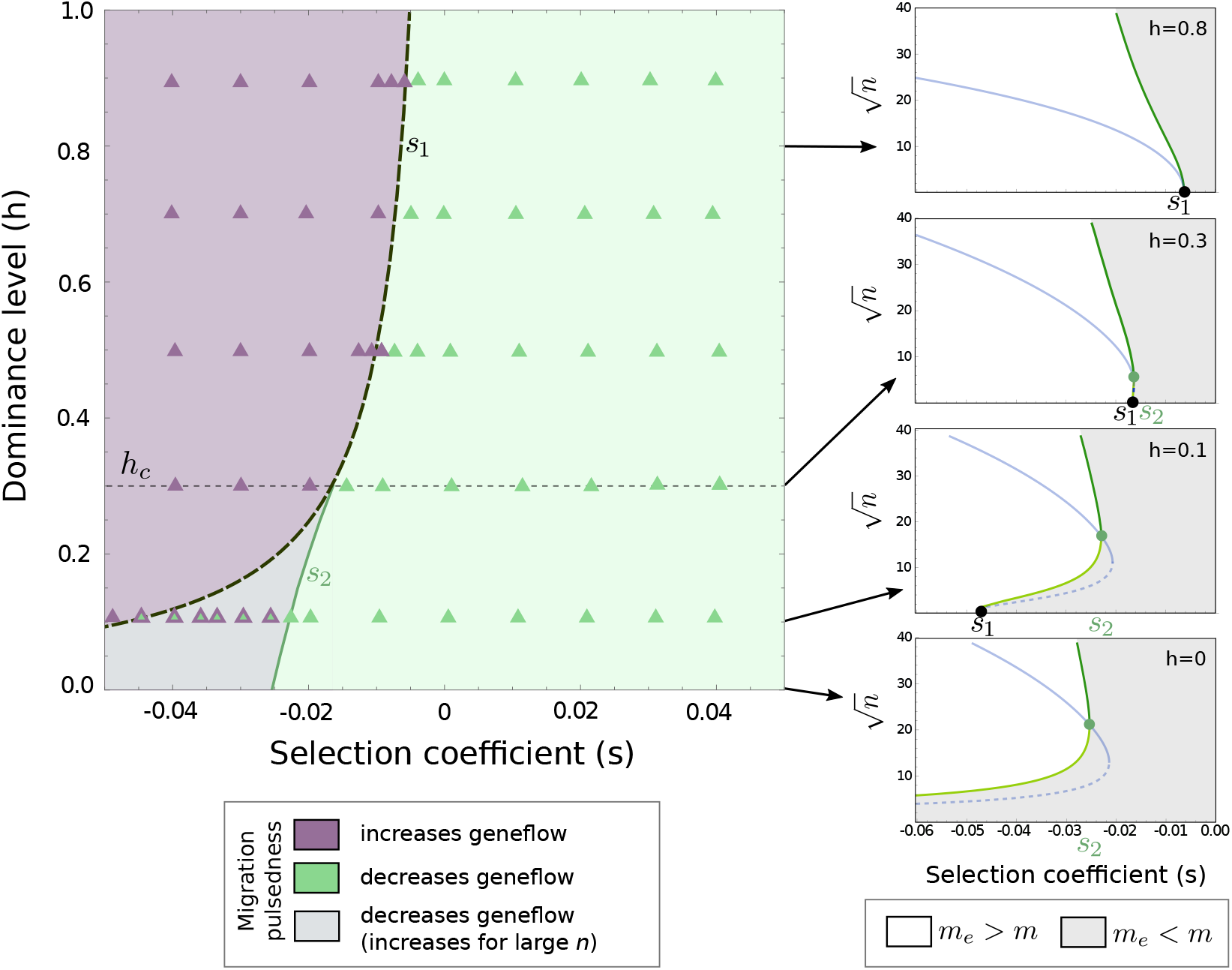
Mathematical predictions. For all selection coefficient *s* and dominance *h* combinations, the plane is partitioned according to the impact of migration pulsedness on effective migration rate (geneflow). Predictions were computed numerically (from eq. (1) and (3)). Triangles show the observed result in stochastic simulations under the very low migration scenario (*m* = 0.001), to check consistency with mathematical analysis. The dashed curve is the mathematical expression for *s*_1_ (eq. (4))). Subpanels are cross-sections of the main panel at four levels of dominance (*h* = 0.8, 0.3, 0.1 and 0), showing *n* values that increase (white) or decrease (gray) the effective migration rate. Note the square-root y-scale. The blue curves show the *n* values that maximize (solid) or minimize (dotted) geneflow. Other parameters: *N* = *N_e_* = 100.

Differentiating *u*[*f*(*n*)] twice at *n* = 1, using eq. (3) and *f* = *n*/(*N* + *n*), we find after some calculations (S.I.; section 2.3) that migration pulsedness increases the effective migration rate (promotes geneflow) if:

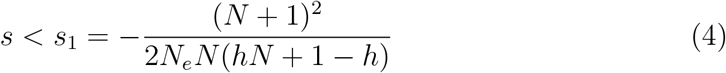

This threshold selection coefficient is always negative. In the special case of additivity (*h* = 1/2), if population size is large and no different from the effective population size, we obtain the approximation *s*_1_ ≈ –1/*N*. This becomes *s*_1_ ≈ –1/2*N* for fully dominant alleles, whereas for fully recessive alleles, we get the quite extreme value *s*_1_ ≈ –1/2. These results are in close agreement with the numerics and with stochastic simulations under a very low migration scenario (fig. 2).

We remark that the values of *n* at which an initially positive effect of pulsedness would revert to negative are usually unrealistically large (thick green curves in sub-panels of fig. 2; see also S.I.; section 2.6). Therefore, eq. (4) in practice means that for sufficiently deleterious alleles (*s* < *s*_1_), migration pulsedness increases the effective migration rate, instead of reducing it as it does for neutral alleles. It reduces geneflow, as for neutral alleles, for slightly deleterious alleles (*s*_1_ < *s* < 0), and for all beneficial alleles (*s* > 0).

The condition *s* < *s*_1_ is sufficient, but there still is the possibility that function *u*(*n*) is first convex, then switches to concave to such an extent that criterion (1) is met for some larger *n* value, even though it was not for small values. In such circumstances, the consequences of migration pulsedness further depend on the value of *n*: it would generally decrease the effective migration rate, except for some intermediate range of *n* values, for which it would on the contrary increase it. This situation requires function *u*(*f*) to have a pronounced sigmoidal shape, which is known to occur only for recessive deleterious alleles.

Accordingly, as can be seen in fig. 2, for sufficiently recessive alleles, increasing *s* above *s*_1_ may not immediately turn the effects of migration pulsedness from positive to negative, but rather there can be an intermediate range (*s*_1_, *s*_2_) of values for which migration pulsedness has mixed effects depending on its intensity. One can determine the level of dominance at which this bifurcation occurs (*h_c_*), by remarking it is the point (*h, s*_1_(*h*)) at which the second inflexion point of function *u*(*n*) collides with the first one (located at *n* = 1). In other words, it is the point at which function *u*(*n*) has a null third derivative at *n* = 1 while verifying the condition *s* = *s*_1_ (S.I.; section 2.4). After some calculations, we get the following condition:

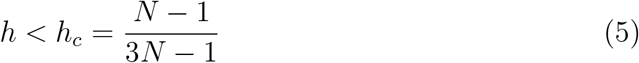

Note that this depends on the census population only, not on the effective population size. If population size is large, we get the approximation *h* ≈ 1/3, in close agreement with the numerics (fig. 2).

We remark that the minimal level of pulsedness required to get an increased effective migration rate is typically quite large (40–100; thin green curves in the subpanels of fig. 2). Therefore, in practice, this situation is essentially similar to that of neutral alleles (migration pulsedness reduces the effective migration rate).

To summarize, migration pulsedness should increase the effective migration rate for sufficiently maladapted and not-too-recessive alleles (*s* < *s*_1_), whereas it should reduce the effective migration rate for slightly maladapted alleles, neutral alleles and beneficial alleles.

### Confronting predictions with stochastic simulations

The above mathematical predictions were well verified in stochastic simulations in which the overall rate of migration *m* was allowed to take larger values (fig. 3). To facilitate comparisons across parameter sets, we follow the usual practice of plotting results in terms of *N_e_s* rather than *s*, to reflect the effective strength of selection relative to drift (Wright, 1931; Lande, 1994).

**Fig. 3.**
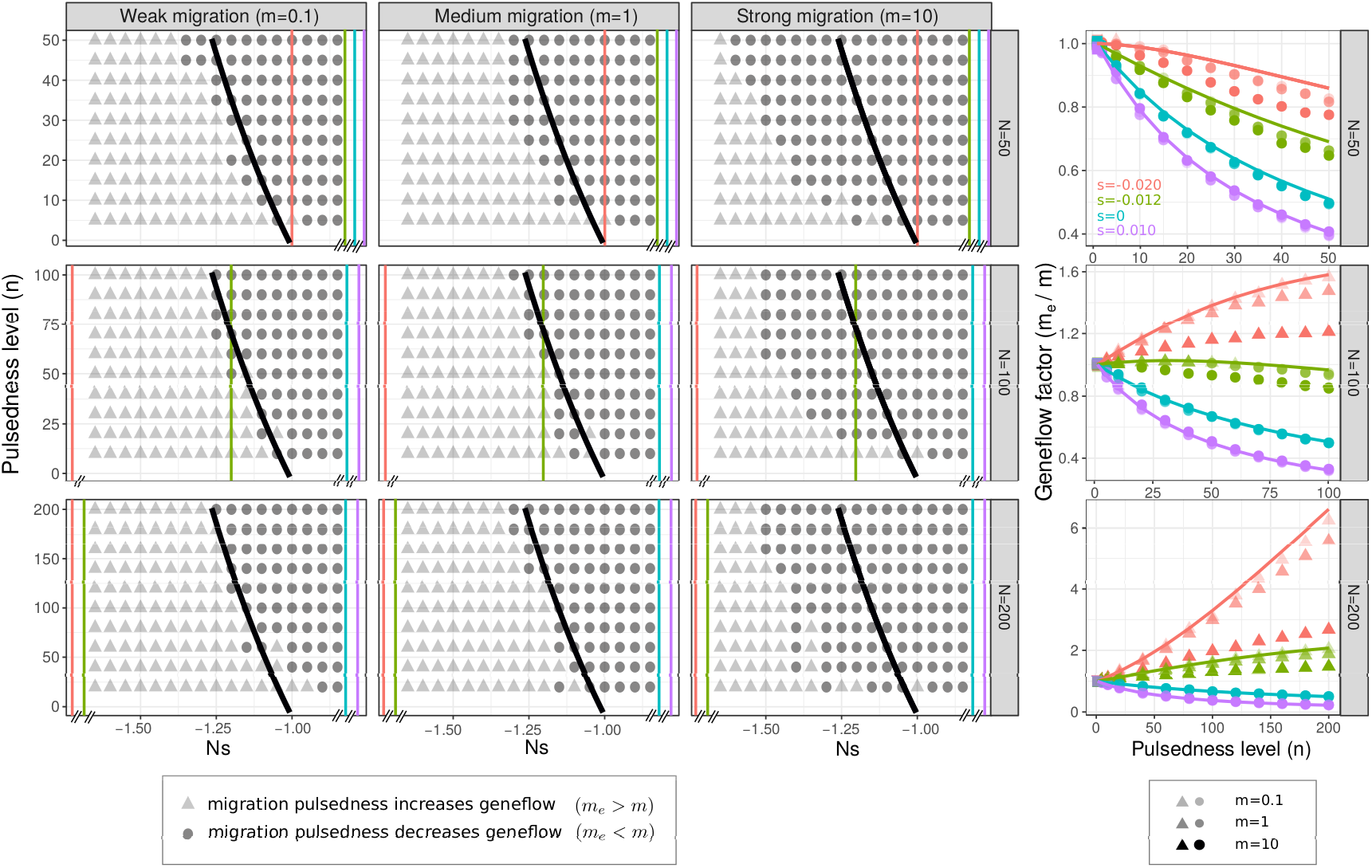
Results of stochastic simulations. *Left:* Effect of migration pulsedness on effective migration rate, for *n* between 1 and *N*, and for a range of *Ns* values. The solid black lines represent the mathematical prediction, obtained numerically from eq. (1) and (3). Results are provided for the tree scenarios of migration intensity (columns), and for three population sizes (rows). *Right*: Geneflow factor (*m_e_/m*) as a function of *n*, for different selection coefficients *s* and for the three migration scenarios. The *s* values used are also positioned in the left panels as vertical lines. Symbols are the values obtained in stochastic simulations, and solid lines are mathematical predictions, obtained as before. Other parameters: *h* = 0.5.

Qualitatively, mathematical predictions hold almost perfectly in the low and medium migration scenarios. In other words, migration pulsedness could either increase or decrease the effective migration rate, depending on the type of selective regime, in the direction predicted. In the strong migration scenario, predictions also held except that the space of selective regimes for which migration pulsedness increases the effective migration rate was shifted to lower *s* values (i.e. *s*_1_ gets more negative).

Results did not depend importantly on population size, provided one scales *n* with *N*. This is again consistent with mathematical predictions: the left-hand side of eq. 1 is unchanged if *N_e_s* and *n/N* (and thus *f*) are both kept constant (see eq. 3). The right-hand side changes very little: its relative variation with *N* is of order less than 1/*N*^2^ (see S.I.; section 2.7). Therefore, mathematical predictions are almost invariant for given values of *N_e_s* and *n/N*.

In quantitative terms, the impact of migration pulsedness on the effective migration rate (the geneflow factor) was very close to the one predicted by our mathematical approximation in all cases, except for deleterious alleles under the strong migration scenario (fig. 3). In the latter case, the observed increase in geneflow could be much smaller than the one predicted. In the most extreme example, a predicted six-fold increase in effective migration rate turned out to be slightly more than a two-fold increase with *m* = 10 (fig. 3). Overall, large migration rates (i.e. beyond the one-migrant-per-generation rule) shift predictions towards a decrease in effective migration rate: more negative *s* values are required to obtain the same impact as for low migration rates. They otherwise have little qualitative impact. Remark that these quantitative differences are most important for large pulsedness levels (*n*); results for low *n* values, and thus the value of *s*_1_, are in much closer agreement with mathematical predictions (fig. 3). Similar conclusions hold for other values of the dominance level *h* (see S.I.; fig. S1).

### Genome-wide consequences: mean fitness and the pulsedness load

The above results have established that migration pulsedness should affect different loci in different ways, depending on their selective regime. Overall, it will promote geneflow (promote fixation) of deleterious or maladapted alleles, especially if not too-recessive, whereas it will reduce geneflow (slowdown fixation) of slightly deleterious, neutral and beneficial alleles, especially recessive ones.

As pulsedness promotes the fixation of maladapted alleles and opposes the fixation of beneficial alleles, its overall effect is to homogenize geneflow (fixation rates) across loci with different selective regimes in a genome. Such a homogeneizing effect could at first be regarded as similar to a decrease in effective population size: demo-graphic fluctuations or other sources of variability that decrease *N_e_* would reduce the strength of selection and also homogenize fixation rates across different loci.

However, simply decreasing population size, in a model with continuous migration, cannot reproduce the patterns observed with migration pulsedness (fig. 4). Even if adjusting potentially unknown parameters (*m* or the time since onset of migration), in order to match exactly the fixation rate of neutral alleles, a reduction in *N_e_* can either fit the fixation patterns observed for maladapted alleles, or those of beneficial alleles, but never fits both (fig. 4). In other words, the change in *N_e_* that is required to mimic the consequences of migration pulsedness is not identical across loci differing in their selection regimes. Therefore, the consequences of migration pulsedness cannot be interpreted as some reduction in effective population size.

**Fig. 4.**
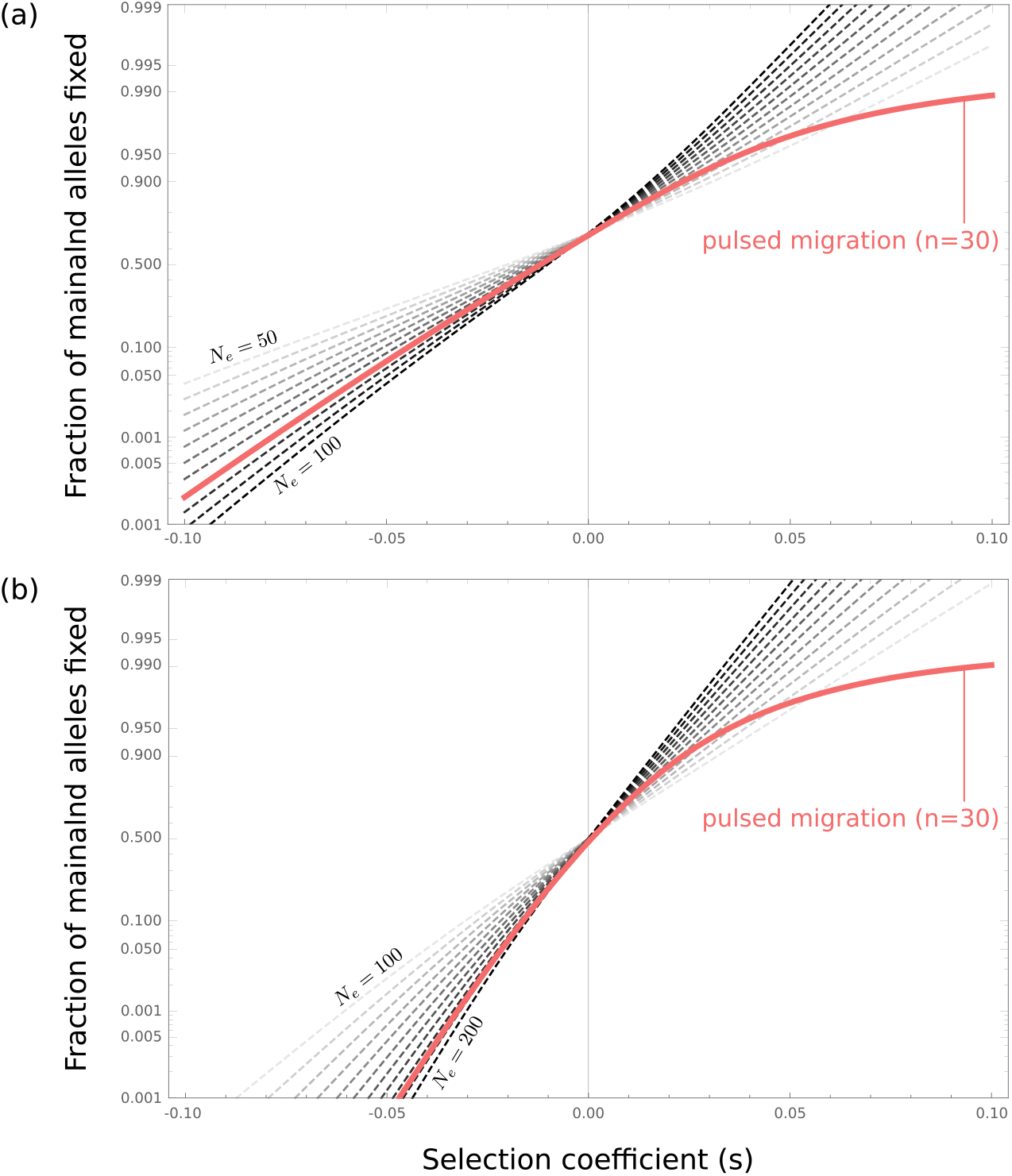
Probability of fixation of different alleles varying in their selection coefficient, under pulsed migration (*n* = 30; red curve), and under continuous migration with different effective population sizes (gray dashed curves). (a) *N* =100 and (b) *N* = 200. The time since the onset of migration (*t*) was set at an intermediate value so that about 50% of neutral alleles have gone to fixation in the pulsed case: *t* = 150. To minimize the difference between pulsed and continuous curves, the time was adjusted for the continuous cases, so as to match the neutral fixation rate: (a) *t* = 120 and (b) *t* = 140. If using the actual times, the discrepancies between the two migration scenarios are even more pronounced. The curves are from the mathematical model with parameters *m* = 1 and *h* = 1/2. Note the logistic scale for the y-axis.

One can extrapolate the fitness consequences of migration pulsedness over the entire genome, assuming a large number of independent loci, each with its particular selection regime. The distribution of *h* and *s* values over loci would vary greatly across organisms and contexts, but for illustration we can assume some plausible distribution of fitness effects (DFE) with *h* = 1/2, such as the one proposed in (Martin & Lenormand, 2015, S.I.; section 3.1). This DFE, illustrated in fig. 5a, features an excess of deleterious (maladapted) alleles, representative of situations where migration from the mainland mostly decreases fitness on the island. At any time, the mean fitness 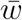 in the island population is the product of the fitness contributions of all loci, depending on their fixation status (see S.I.; section 3.2).

**Fig. 5.**
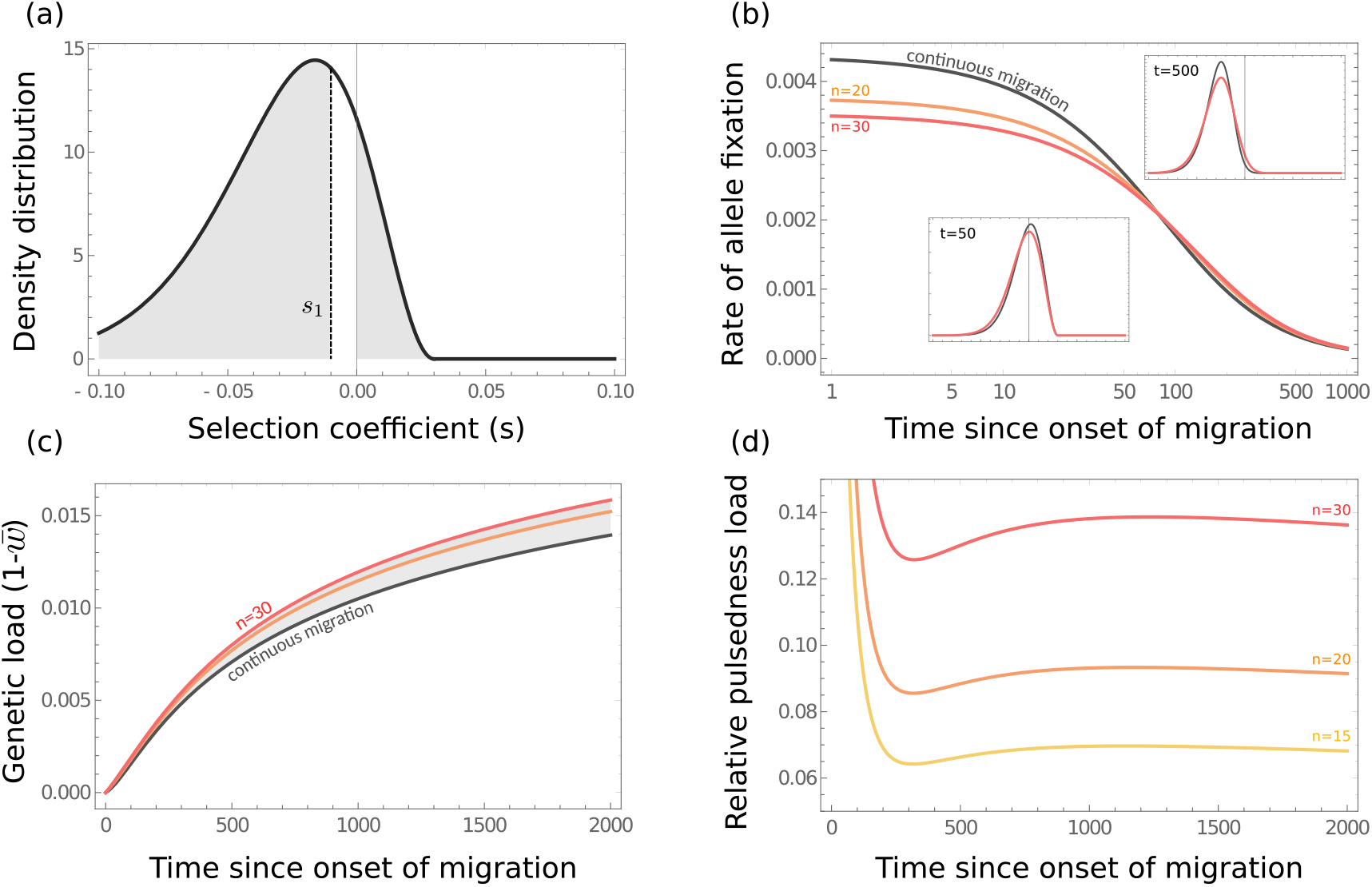
The pulsedness load. (a) An illustrative distribution of fitness effects (DFE) for mainland alleles across loci. Most alleles are slightly deleterious or neutral, some are beneficial (Martin & Lenormand, 2015). Three important classes of alleles are distinguished: *s* < *s*_1_, *s*_1_ < *s* < 0 and *s* > 0. (b) The instantaneous rate of fixation (fixation flux) of mainland alleles as a function of the time since onset of migration, for continuous migration and two levels of migration pulsedness. The two inserts show the DFE of alleles that are currently going to fixation, at times 50 and 500, for continuous migration and pulsed (*n* = 20 and *n* = 30) migration. (c) Genetic load (1– mean fitness) as a function of time since onset of migration, for continuous and pulsed migration. The excess of genetic load caused by migration pulsedness (the pulsed load) is highlighted as the shaded area. (d) Value of the pulsedness load (relative to the genetic load with continuous migration) as a function of time since onset of migration, for three different levels of migration pulsedness. Results are from the mathematical model with parameters *h* =1/2, *m* = 1 and *N* = 100.

Following the onset of immigration, the island population mean fitness evolves through time, asymptotically settling at its equilibrium value, when all mainland alleles have fixed, and thus controlled by the DFE. Under recurring migration, regardless the level of pulsedness, beneficial (adapted) alleles tend to go to fixation first, followed by neutral alleles, and ultimately deleterious (maladapted) alleles. These dynamics of allele fixation are shown in fig. 5b, with the FDE of fixating alleles, at two different times, is shown as inserts. Introducing migration pulsedness slows down the fixation of beneficial and neutral variation, which decreases mean fitness in the island population, compared to what it should be under continuous migration. Conversely, it hastens the fixation of deleterious (maladapted) alleles with *s* < *s*_1_, which is also detrimental to mean fitness (see fig. 5a). As a result, migration pulsedness causes mean fitness to be lower than under continuous migration (fig. 5c), and we call this fitness deficit (i.e. additional genetic load) the “pulsedness load”. If most fitness variation is driven by adaptation to local habitat conditions, the pulsedness load translates into lower local adaptation. It must be stressed that if one simply extrapolated the fact that migration pulsedness decreases the effective migration rate, as is observed for neutral alleles, to the entire genome, one would reach the opposite prediction: migration pulsedness would *increase* mean fitness at any time.

As shown in fig. 5d, the pulsedness load can readily cause a 10% increase in the genetic load, compared to what is under continuous migration. Interestingly, slightly deleterious alleles with selective values in the range (*s*_1_, 0) have reduced fixation rates under pulsed migration, which has the opposite effect of boosting mean fitness: this contributes negatively to the pulsedness load. As a consequence, the pulsedness load might transiently decline (and possibly become negative), at the time when most alleles getting to fixation fall into this category (fig. 5d). Generically, the dynamics of the pulsedness load thus follows three consecutive phases over time, driven by the fixation of different categories of alleles (fig. 5a). The relative importance and timing of the three phases of course depends on the exact DFE.

## Discussion

Temporal variability in migration is probably pervasive in nature. Plants, fungi, birds, mollusks and other marine invertebrates are most represented in the literature about variable dispersal, with many other taxa presumably concerned, including hominoids (Folinsbee & Brooks, 2007). A common form of migration temporal variability is migration pulsedness, i.e. the positively correlated migration of individuals, producing pulses of migration interspersed among periods of low or absent migration. However, classical evolutionary theory largely rests on the assumption of constant migration rates (e.g. Johnson *et al.*, 2000; Yeaman & Otto, 2011; Mailund *et al.*, 2012; Rousset, 2013; Peniston *et al.*, 2019). Considering the growing evidence for non-constant migration processes, it is important to gain a general theoretical understanding of their evolutionary consequences.

We here proposed a novel approach and derived mathematical predictions regarding how the level of migration pulsedness should impact geneflow (the effective migration rate) at loci subject to an arbitrary selection regime. This is an important difference from earlier existing studies, that usually focused on neutral variation alone, or on some specific forms of selection. Though we here addressed pulsed migration patterns (Yamaguchi & Iwasa, 2013; Peniston *et al.*, 2019), we can reasonably expect our predictions to apply more broadly to other forms of variable migration patterns, intended as a temporal overdispersion in the number of migrants per generation. The latter is quantified by parameter *n* in our model.

Our main finding is that the effect of migration pulsedness on geneflow depends not only quantitatively, but also qualitatively, on the selective regime considered. We find that migration pulsedness should decrease the effective migration rate (reduce geneflow) for alleles that are neutral or beneficial. However, for sufficiently deleterious (maladapted) alleles, the effect can be opposite. We found that there exists some threshold selection coefficient *s*_1_, of which we derived a mathematical expression, below which migration pulsedness on the contrary increases the effective migration rate (increases geneflow). The value of *s*_1_ increases with the dominance level *h*, so that an increase in geneflow with pulsedness is more likely for dominant alleles, and much less likely for recessive alleles.

The effect of migration pulsedness is therefore not homogeneous across different loci over the genome that experience different selection regimes. Migration pulsedness increases geneflow, and thus increases the speed and probability of fixation, for deleterious (maladapted) alleles, but it does the opposite for beneficial alleles. This homogeneizes fixation rates over the genome. Such a homogeneization of fixation rates across loci experiencing contrasted selection regimes could also be brought about by a reduced effective population size, i.e. a greater importance of drift relative to selection. However, we showed that a simple change in effective population size, retaining a continuous mode of migration, cannot adequately reproduce the fixation patterns created by pulsed migration. The action of migration pulsedness therefore leaves a signature that, at least in principle, could be distinguished from other forms of variability that reduce effective population size (random population fluctuations, random inter-individual fecundity variations, biased sex-ratios). The signature is more subtle. It is more comparable to that of deterministic processes such as directional changes of population size through time (Otto & Whitlock, 1997). In particular, an increased fixation of deleterious alleles and concomitant decreased fixation of beneficial alleles is analogous of what is predicted for shrinking populations (Otto & Whitlock, 1997). Our results also suggest that migration pulsedness would generate a relative excess of dominant deleterious alleles, and a relative deficit of recessive deleterious alleles (see fig. 2), which adds an extra dimension to its genomic signature.

Another consequence of migration pulsedness is that, by promoting the fixation of deleterious (maladapted) alleles, while slowing down the fixation of beneficial (adaptive) alleles, it overall reduces mean fitness and the level of local adaptation. We showed that migration pulsedness creates an additional genetic load, on top of the migration load expected under a continuous migration of similar intensity. This so-called “pulsedness load” can be non-negligible, and may easily represent a 10% increase in the genetic load, or even more, for plausible distributions of fitness effects and pulsedness values.

Our results generalize some existing results and are compatible with them. Most importantly, our predictions for neutral alleles are consistent with the few earlier population genetics studies, even though they used entirely different modelling approaches (Nagylaki, 1979; Latter & Sved, 1981; Whitlock, 1992; Rousset, 2013). We predict, as they did, that temporal variation in migration rates always decreases the effective migration rate for neutral alleles.

Even fewer studies have so far considered non-neutral variation. Peniston *et al.* (2019) investigated the consequences of pulses of migration in a mainland island model, like us, but they specifically considered source-sink situations: the island population was initially not viable on its own, and had to adapt in the face of maladaptive geneflow. This is quite a different scenario, as we here assumed that the island population was viable, not a sink. Our results thus complement the latter study, and we find, as they did, that migration pulsedness can importantly affect the dynamics of local adaptation. Even though direct comparisons are difficult, we remark that among all the scenarios they considered, for the one closest to ours, Peniston *et al.* (2019) report that more pulsedness migration hampers local adaptation, which is consistent with our predictions.

Yamaguchi & Iwasa (2013) and subsequent papers considered parapatric speciation through the accumulation of neutral mutations between two isolated populations, until prezygotic isolation develops. They used a formalism similar to ours, but were not specifically interested in migration variability. They did not vary the latter, and for simplicity they kept the number of migrants per migration event (*n*) at very low values. Their results thus cannot be compared to ours. However, we can use our results to predict the consequences for the build-up of reproductive isolation between isolated populations. Indeed, if we extend the range of selection regimes we’ve considered to incompatible mutations causing reproductive isolation (Dobzhanski-Muller – DM – mutations; Gavrilets (2004)), we can see those as the limit where *s* → *0*^-^, and *h* → ∞ (strong overdominance). For instance, *s* = −0.001 and *h* = 10000 describes a mutation that cause a −0.1 fitness disadvantage, only in heterozygotes. Extrapolating fig. (2) to those values, we would conclude that migration pulsedness should slightly decrease geneflow for them. This may seem paradoxical, as such mutations obviously benefit from arriving in higher frequency. However, their probability to go to fixation by chance is almost zero, unless their frequency reaches or exceeds 50%, a value that would require unrealistic large values of *n*. We can tentatively posit that migration pulsedness should not much affect the accumulation of DM incompatibilities, even though establishing this would require a dedicated study.

Most of our predictions can be understood in terms of the trade-off between fewer migration events (rate *m/n*), which reduces geneflow, and higher initial frequency after an event (*n*/(*N* + *n*)), which promotes geneflow. Increasing the pulsedness level *n* decreases the number of migration events in inverse proportion, but the initial frequency at each event increases less than linearly (*n*/(*N* + *n*)). In general, and for neutral alleles in particular, the net effect of increasing *n* is therefore detrimental to geneflow. Intuitively, this is because concentrating the arrival of alleles in time causes those mainland alleles to compete more among themselves, which does not help their fixation over local alleles. It is only when the fact of arriving in larger initial frequency is very beneficial to fixation that this effect can overcome that of a decreased number of migration events. This is the case for sufficiently deleterious alleles, whose chance of establishing is smallish from a low initial frequency, but much better if in higher frequency, owing to genetic drift. We have shown that sufficiently deleterious here means *s* < *s*_1_. The beneficial effect of a larger initial frequency is of course strengthened by positive frequency-dependent selection: this is why the value of *s*_1_ increases with the level of dominance *h* in a diploid context, as it generates positive frequency-dependence at the allele level. An increase in geneflow with migration pulsedness would similarly be promoted by other sources of positive frequency-dependence, such as aposematic color signalling (Endler, 1988): migration pulsedness could probably promote the successful migration of foreign aposematic morphs.

Our mathematical predictions held very well in stochastic simulations for different population sizes, and when the overall migration rate (*m*) was large. However, in the latter case the possibility for migration pulsedness to increase geneflow is shifted towards more deleterious alleles, i.e. requires more stringent conditions than at lower migration rates. This can be understood intuitively in terms of mass-effects: with large migration rates, the influx of mainland alleles is strong enough to increase their frequency in the island on its own, regardless of the local selection-drift dynamics. In other words, deleterious alleles “pile-up” over successive migration events, and this can be an important driver of their eventual fixation. The latter effect is virtually unaffected by migration pulsedness; it solely depends on the average rate *m*. As a consequence, the positive impact of migration pulsedness that we have identified, and explained just before, is in some sense diluted and gets relatively weaker. This is not the case for beneficial alleles, which get fixed after a small number of migration events, so that the “piling-up” effect does not significantly contribute to their fixation. Further mathematical investigations allowing for large migration would be needed to validate this interpretation.

The finding that migration pulsedness all else equal favors the fixation of deleterious alleles and reduces local adaptation has direct implications for biological conservation, population management and island biology. For instance, population reinforcement programs usually takes the form of periodic releases of groups of individuals, whose frequency and intensity must both be optimized. Our results suggest that rare introductions of relatively large numbers of individuals, i.e. pulsed introduction patterns, would be more detrimental to population fitness and local adaptation, relative to more continuous fluxes of individuals, possibly compromising population viability and persistence. Extending our reasoning to island communities (Cowie & Holland, 2006), this also suggests that sporadic but potentially intense bouts of immigration, as brought about by rafting and human-driven invasions, could favour the establishment of relatively maladapted mainland species within communities competing for similar resources.

Of course, these impacts would combine with other effects, in particular demo-graphical effects. It has been shown for instance that in small populations subject to Allee effects, pulsed migration patterns can be favourable to population establishment and persistence (Rajakaruna *et al.*, 2013; Bajeux *et al.*, 2019). Similarly, Peniston *et al.* (2019) found that migration pulsedness could sometimes favour local adaptation and persistence in populations that are demographic sinks (see also Gaggiotti & Smouse, 1996). These examples underline how complex the consequences of dispersal can be when demo-genetics effects are taken into account (Garant *et al.*, 2007). It would be interesting to extend our model considering a broader range of demographic scenarios, and see how predictions are affected. For instance, the decrease in mean fitness through time as mainland alleles fix in the island population could have a negative impact on the local population. A decrease in *N* could have various impacts on the probabilities of fixation. At the simplest level, it would increase the *n/N* ratio, which would make the effects of pulsedness more and more pronounced. It would also increase the importance of drift, i.e. decrease *N_e_s*, which would shift the value of *s*_1_. How exactly this would impact the overall dynamics of the pulsedness load remains to be determined. At a more genetic level, we assumed for simplicity that alleles at different loci were unlinked and behaved independently. This would not always be the case, and it would be interesting to study the role of migration pulsedness for pairs of linked mutations that are in strong linkage disequilibrium among migrant individuals.

We conclude, along the lines of Peniston *et al.* (2019), by advocating for tests of predictions on migration pulsedness in empirical systems with well-designed experimental set-ups. We add that available data on migration flows, e.g. from monitoring of oceanic rafting, individual tracking of dispersal routes, or marine and aerial current models (Ser-Giacomi *et al.*, 2015; Lagomarsino Oneto *et al.*, 2020), combined with genomics data, may also provide opportunities for testing such predictions in the wild. Understanding the consequences of ecological variability is increasingly important in the present context of climate and habitat changes, and the spatio-temporal variability in migration patterns probably deserves to receive more attention.

## Supporting information

aubree_et_al_2022_SI.pdf

