## Supplementary material for "Migration pulsedness alters patterns of allele fixation and local adaptation in a mainland-island model": aubree_et_al_2022_SI.pdf

September 30, 2022

**4 Supplementary Figure 1: stochastic simulations with dominance** **11**

### 1 Stochastic simulations

#### 1.1 Gillespie algorithm for the simulations

We perform fully stochastic simulations of the island population partially connected to a mainland population, using a Gillespie algorithm to model a logistic demography of island population with migration (Goel & Richter-Dyn, 1974).

There are three possible events: birth, death, and migration from mainland to island (rate $m/n$), whose rates are reevaluated at every iteration steps and are used to weight the random choice of the next occurring event. To avoid too frequent population extinctions during the simulations, birth rate is taken proportional to the population size $N_t$ at time $t$ while death rate is taken proportional to $\frac{N_t^2}{N}$ (logistic demography). The factor of proportionality $d$ scales the unit time to a given number of generation. A generation lasts $\frac{N}{d}$ units time in average, and $d$ is taken equal to $N$. When death occurs, an individual is randomly drawn and removed from the population. At birth, two random individuals are drawn with replacement, with a probability that depends on their fitness (fertility selection), and one allele of each is randomly chosen to form the new born. Because our approach only models selection at birth and not at death and to make our selection coefficient comparable to the one classically used (Kimura, 1962), we multiplied by 2 the effect of selection felt by individuals at birth. When a migration event occur, $2n$ alleles $A$ immigrate into the island. After event application, time is incremented of a quantity drawn from a Poisson distribution, whose rate is the sum of the three event rates ($N_t + \frac{N_t^2}{N} + \frac{m}{n}$).

The first migration event occurs on average $\frac{n}{m}$ generations after simulation starts. This model the case when populations could be physically connected , but where stochasticity may not have yet allowed individuals to move from one to another. We chose to not set time start to the first migration event, because this would diminish the effect that large pulse occur less often (see Bajeux *et al.*, 2019 for a similar simulation start, or also Peniston *et al.*, 2019).

#### 1.2 Optimized algorithm

When migration is rare compared to demographic events, simulation can take a very long time to run (waiting for fixation to occur). To reduce simulation time, we do not simulate demographic events if the island population becomes monomorphic (namely when stochastic birth or death will no longer impact genetic drift). In order not to distort time, we must take into account the time that would have elapsed if we had let the demography unfold.

For this, we increment time by a value drawn into the distribution of migration event times ($\tau_m \, \mathrm{e}^{-\tau_m x}$, with $\tau_m = \frac{m}{n}$ the migration event rate), that is truncated from $\Delta_t$ to infinity. $\Delta_t$ is the elapsed time between the last migration event and the point at which we stopped demography simulation.

This optimization is most effective for simulating low migration intensities ($m = 0.001$ and $m = 0.1$), since long intervals between successive migratione events can be skipped through quickly. Those simulations are also the longest, hence the benefits of this optimization. For large migration values, the optimization has practically no effect, and amounts to using the basic Gillespie algorithm.

#### 1.3 Obtaining the effective migration rate from the simulations

Effective migration rates are calculated from the fixation time (time from the onset of migration to the fixation of the mainland allele in the island), using an abacus which gives $m_e$ as a function of fixation time.

To get the abacus, we record fixation times $t_f$ in simulation for $n = 1$ (constant migration) and a large range of migration rates $m$. We then fit the relation between $\log(m)$ and $\log(t_f)$

by a polynomial of order 3 function, which allows to make the correspondence in between any measured fixation time (for any $n$ and $m$) and an effective migration rate. Figure 1 gives an example of such an abacus.

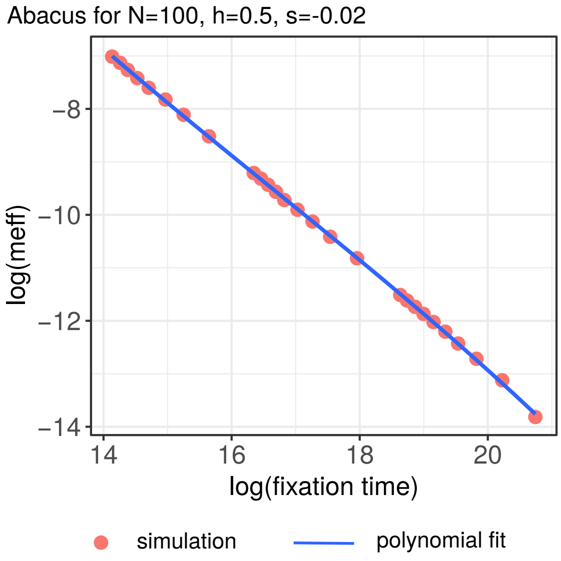

Figure 1: Example of abacus with a polynomial order 3 function fit. For any fixation time obtained in a simulation with the same parameters $N$, $s$ and $h$, we can derive the corresponding effective migration rate.

#### 1.4 Mean number of migration events before fixation

We recorded the mean number of migration events before fixation occurs in the simulation (fig. 2). We can verify that that his number does decline as $1/n$, as expected mathematically, and that in most cases, even for very pulsed migration scenarios, at least three events are required for fixation to occur, even for neutral alleles. Obvisouly, larger values of $m$ tend to decrease these numbers, especially for counterselected alleles, as the mass effect of migration can by itself drive fixation, regardless of local selection/drift dynamics.

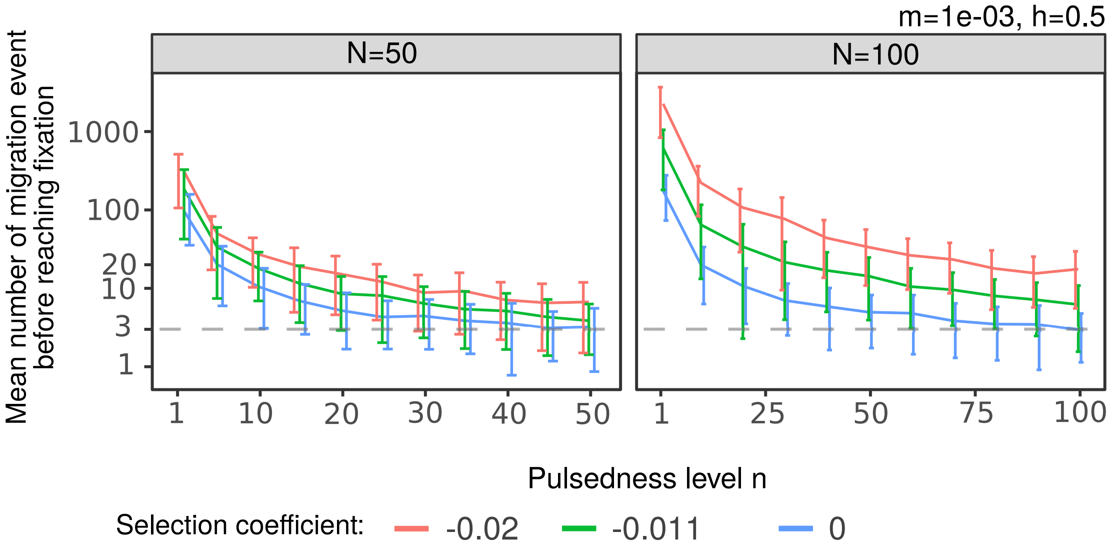

Figure 2: Mean number of migration event before reaching fixation of the mainland allele into the island in log scale, as a function of pulsedness level. Error bar stands for standard deviation.

### 2 Mathematical analysis

#### 2.1 More on the time-scale separation

The time scale separation needed for the mathematical analysis is verified if the average time between two migration events $t_m = \frac{n}{m}$ is large enough so that $t_m \gg t_{c,fix}$, with $t_{c,fix}$ the averaged fixation time before fixation (after a unique initial introduction of an allele into a population). According to Otto & Whitlock (2013), this time is larger for neutral alleles. The general expression of fixation time for any initial allele frequency $f$ and any selection or dominance coefficient can be found in Kimura & Ohta (1969); Whitlock (2003).

At most, $t_{c,fix}$ is the fixation time of one initial copy of the allele of interest in a large population ($f \ll 1$). In that case, $t_{c,fix} \approx 4N$ generations (see also Kimura & Ohta, 1969). On the other hand, the smallest average value of $t_m$ is $1/m$, i.e; for continuous migration. Thus, the time scale separation is verified if $1/m \gg 4NT$, namely for $mT \ll \frac{1}{4N}$, with $T$ the duration of one generation. This is, obviously, a sufficient condition. For non-neutral alleles (that go to fixation or disappear more quickly on average) and for pulsed migration (for which $t_m$ is longer), the condition becomes less stringent, and so the approximation applies more broadly. It is thus a useful approximation as these are the situations we are most interested in here.

#### 2.2 Probability $f_1$ that a mainland allele has fixed

For a given selection value $s$ and dominance level $h$, the probability that the mainland allele has fixed in the island at time $t$, $f_1(t)$, changes as:

$$\frac{df_1}{dt} = \beta(1 - f_1)$$

with $\beta = \frac{m}{n}u(f, N_e, s, h)$ and $u(f, N_e, s, h)$ the fixation probability of an allele arriving in frequency $f = \frac{n}{n+N}$. Solving this differential equation leaves us with

$$
\begin{aligned}
f_1(t) &= 1 - (1 - f_1(0)) \exp(-\beta t) \\
&= 1 - (1 - f_1(0)) \exp\left[-\frac{m}{n} u\left(\frac{n}{n+N}, N_e, s, h\right) t\right]
\end{aligned} \tag{1}
$$

We'll assume that all mainland alleles are initially absent from the island population, i.e. we use the initial condition $f_1(0) = 0$. In this context, the mean time to fixation ($t_f$) is $1/\beta$.

#### 2.3 Derivation of the value of $s_1$

**Preliminary lemma: initial tangientiality** Let us differentiate the two sides of the general citerion (1) w.r.t $n$, at $n = 1$, and then assumie that $N$ is large, i.e. we $N \gg 1$ and neglect terms $\mathcal{O}(1/N^2)$. After some calculations we get for the left hand side:

$$\frac{N - 4N_e hs}{N^2(1 - 2N_e hs)}$$

and for the right hand side:

$$\frac{N - 2N_e hs}{N^2(1 - 2N_e hs)}$$

The ratio of the two slopes is therefore

$$2 - \frac{N}{N - 2N_e hs}$$

We remark that for weak selection, more specifically for $hs$ close to zero, and more so if $N_e < N$, the above ratio is almost equal to one. From numerical investigations (Fig. 2), these conditions are always verified when we are near the threshold value $s_1$. In other words, to a good approximation the two sides of the general criterion have at $n = 1$ the same value and the same slope with respect to $n$.

Therefore, the general criterion (1) can be seen as a condition on the curvature of function $u(n)$ at $n = 1$: if the curvature is positive, the criterion is met, if the curvature is negative, the criterion is not met.

**The value of $s_1$** We differentiate $u(n)$ (eq. (3) in the main text) twice with respect to $n$, and evaluate it at $n = 1$. We get:

$$\frac{f'(1)G'(f(1)) + G(f(1))f''(1)}{(N+1)^4}$$

where $f(n) = n/(N+n)$ and $G(f)$ is as defined in the main text. For clarity, prime symbols are used for differentiation, i.e. prime denotes differentiation w.r.t. $n$ for $f$, and differentiation w.r.t. $f$ for $G$.

Evaluating the above expression, and solving for zero in terms of $s$ (intermediate steps are omitted), we obtain:

$$s_1 = \frac{(N+1)^2}{2NN_e(1 - h + hN)}$$

which the value reported in the main article.

The approximations given in the main text follow from further assuming $N = N_e$ and ignoring terms $\mathcal{O}(1/N^2)$.

#### 2.4 Derivation of the value of $h_c$

The value of $h_c$ is obtained as the value such as the curvature of $u(n)$ is null (for we are at $s = s_1$) and does not change with $n$. In other words, the third-order derivative of $u(n)$ with respect to $n$ vanishes.

The third-derivative of $u(n)$ is

$$\frac{3f'(1)G'(f(1))f''(1) + f''(1)^3 G''(f(1)) + G(f(1))f'''(1)}{(N+1)^6}$$

As before, we compute the above expression by performing the necessary differentiations, and we evaluate it at $s = s_1$ using the above derived value. We deduce that the thrid derivative cancels out if

$$\frac{2N(N+1)^4(1 + h(3N-1) - N)}{1 + h(N-1)} = 0$$

Finally, solving for $h$ we get

$$h_C = \frac{N-1}{3N-1}$$

which is the value reported in the main article. The approximation follows directly from $N \ll 1$.

#### 2.5 Derivation of the slope of $n_+$ around $s_1$ as a function of $s$

There exists a value $n_+$ of $n$ that maximises the difference in fixation rate in between the continuous case and the pulsed case. Owing to the fact that fixation rate is proportional to the ratio $\frac{u(s,n)}{n}$, we can study the latter to find $n_+$. Along a curve $(s, n_+(s))$ the following condition is verified:

$$\left.\frac{\partial}{\partial n}\frac{u(s,n)}{n}\right|_{(s,n_+(s))} = 0 \tag{2}$$

Similarly to the previous paragraph, we differentiate equation (2) with respect to $s$. We use the notation used in the previous paragraph.

$$\frac{d}{ds}\left(\left.\frac{\partial}{\partial n}\frac{u(s,n)}{n}\right|_{(s,n_+(s))}\right)(s, n_+(s)) = 0$$

with

$$\begin{aligned}\frac{d}{ds}\left(\left.\frac{\partial}{\partial n}\frac{u(s,n)}{n}\right|_{(s,n_+(s))}\right)(s, n_+(s)) =& \frac{2u(s,n_+(s))\frac{dn_+}{ds}}{n_+^3(s)} \\ &- \frac{2\frac{dn_+}{ds}u^n(s,n_+(s)) + u^s(s,n_+(s))}{n_+^2(s)} \\ &+ \frac{\frac{dn_+}{ds}u^{n^2}(s,n_+(s)) + u^{s,n}(s,n_+(s))}{n_+(s)}\end{aligned}$$

Then we isolate the slope $\frac{dn_+}{ds}$ and find:

$$\frac{dn_+}{ds} = \frac{n_+(s)\left(n_+(s)u^{s,n}(s,n_+(s)) - u^s(s,n_+(s))\right)}{2u(s,n_+(s)) - 2n_+(s)u^n(s,n_+(s)) + n_+^2(s)u^{n^2}(s,n_+(s))}$$

Such as in the previous case, at this stage, the expression of the slope $\frac{dn_+}{ds}$ is valid for any shape of $u$. We can express this slope in the particular case of frequency independent selection where $u = \frac{1-\mathrm{e}^{-2Ns\frac{n}{n+N}}}{1-\mathrm{e}^{-2Ns}}$. In that case, we can use the above approximation $(s_1, n_+(s_1)) \simeq (-1/N, 1)$, so we can know the slope at $s_1$. A Taylor expansion for large $N$ ($N \gg 1$) gives:

$$\left.\frac{dn_+}{ds}\right|_{s=s_1} \simeq -\frac{3}{2}N^2 \tag{3}$$

This result is well verified numerically (see fig. **??**). We thus found that the slope of $n_+$ as a function of $s$ close to the limit $s_1$ is well approximated by $-\frac{3}{2}N^2$.

#### 2.6 Derivation of the slope of $n_l$, knowing the slope of $n_+$

Along a curve $(s, n_l(s))$ the following condition is verified:

$$u(s, n_l(s)) = n_l(s)u(s, 1) \tag{4}$$

Solving this condition, we numerically observe that the slope of $n_l$ close to $s_1$ is approximated by $-\frac{3}{2}N^2$ (see fig. **??**). It is twice the slope found for $n_+$ : $\left.\frac{dn_+}{ds}\right|_{s=s_1} \simeq -\frac{3}{4}N^2$ (see previous paragraph). It means that, for a given $s$ close to $s_{l1}$, the pusledness value $n = 2n_+$ should verify the condition for the effect of pulsedness, i.e.

$$u(s, 2n_+(s)) = 2 * n_+(s)u(s, 1) \tag{5}$$

Let's verify this relationship. We note $\Delta_c(s) = u(s, 2 * n_+(s)) - 2 * n_+(s)u(s, 1)$, and we want to verify whether or not it cancels close to $s_{l1}$. We know that $(s_1, n_+(s_{l1})) \simeq (-2/N, 1)$. Thus, close to $s_1$, the equation of function $n_+(s)$ is

$$n_+(s) = -\frac{3N^4}{2}s - \frac{3N}{2}$$

Replacing this expression into $\Delta_c$ gives:

$$\Delta_c(s) = \left(1 + \frac{1}{-1 + \mathrm{e}^{Ns}}\right)\left(1 - \mathrm{e}^{-\frac{3Ns(2+Ns)}{4+3Ns}} + 3N + \frac{3}{2}N\left(Ns - \mathrm{e}^{-\frac{Ns}{1+N}}(2 + Ns)\right)\right)$$

We then evaluate $\Delta_c$ at $s = s_1 + ds = -\frac{2}{N} + ds$, and look for the Taylor expansion for $ds \ll 1$ and large $N \gg 1$. Neglecting the terms $\mathcal{O}(ds^2)$, we obtain that

$$\Delta_c(s_{l1} + ds) \simeq \frac{2}{(\mathrm{e}^2 - 1)N}ds$$

Then

$$\lim_{ds \to 0} \Delta_c(s_{l1} + ds) = 0$$

Note that this limit was already true without the Taylor expansion for small $ds$. This limit means that close to $s_{l1}$, the condition (5) is verified.

We thus have shown that

$$n_l = 2n_+$$

close to $s_1$, and the slope of $n_l$ as a function of $s$ is twice the slope of $n_+$ (see fig. **??**):

$$\left.\frac{dn_l}{ds}\right|_{s=s_1} \simeq -2\frac{3}{2}N^2 \simeq -3N^2 \tag{6}$$

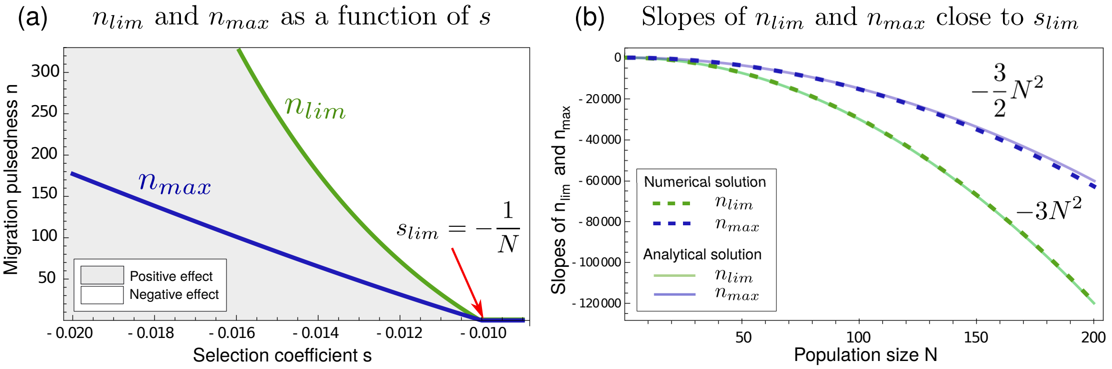

Figure 3: (a) $n_l$ and $n_+$ as a function of $s$, found numerically solving equation (4) for $n_l$ and equation (2) for $n_+$. The limit value $s_{l1}$ of selection bellow which the qualitative effect of migration pulsedness depends on the value of $n$ is indicated with a red arrow. It has been analytically estimated to be equal to $s_{l1} = -\frac{1}{N}$. Here $N = 100$. (b) Close to this selection limit value, the slopes of $n_l$ and $n_+$ are estimated both numerically (doted lines) and analytically (solid lines).

#### 2.7 Invariance of the criterion

Let us consider that the ratio $n/(N+n)$ is kept unchanged upon changing $N$, i.e. $n = aN$. Let us also assume that effective population size scales with census population size, i.e. $N_e = bN$. It follows that if we keep the $N_e s$ ratio constant, we have $s = c/N_e = c/(bN)$. We first observe that function $u(n)$ is invariant in this case, since function $G$ is unchanged if $N_e s$ is constant, and the inegration bound $f$ is also unchanged (eq. (3) in the main document). Therefore, the left hand side of the criterion is unchanged.

Second, we compute the derivative of the right hand side with respect to $N$, under the above contraints, which yields:

$$\frac{u(1/(N+1))}{N}$$

For weak seleection, $s$ is close to zero, and $u$ can be approximated as its value in the neutral case (eq. (2) in the main text), yielding:

$$\frac{1}{N+N^2}$$

which is very small for even reasonably large $N$.

For $s < 0$, the derivative will be even smaller than that. Deviations will be larger for positive $s$ values, but this is a case where predictions are always quantitatively the same.

This shows that criterion (1) is almost unchanged upon variations of the population size, provided we scale $n$ and $s$ with the latter.

### 3 Genome-wide predictions and the pulsedness load

#### 3.1 Distribution of fitness effects (DFE)

For illustration purposes, we use the DFE proposed by Martin & Lenormand (2015), based on a non-centered $\chi^2$ distribution, so that the probability density of selective effect $s$ is expressed as:

$$p(s) = \frac{2}{\lambda} X^2 \left( df, \frac{2(\epsilon\mu)}{\lambda}, \frac{2(\epsilon\mu - s)}{\lambda} \right) \tag{7}$$

where $X^2(a, b, c)$ is the probability density of value $c$ from a $\chi^2$ distribution with $a$ degrees of freedom and non-centrality parameter $b$.

Following Martin & Lenormand (2015), we used parameter values $df = 6$, $\epsilon = 3$, $\mu = 0.01$ and $\lambda = 0.01$, top roduce the distribution shown in fig. 5 of the main article. THis produces a distribution with reasonable width and clearly biased toward maladapted alleles. Other DFEs give similar conclusions, though results would differ quantitatively, since the relative frequency of the three classes of alleles would vary, as explained in the main text.

For simplicity, additivity is otherwise assumed ($h = 1/2$). Of course, everything that follows is straightforward to extend to the full bi-dimensional case $(s, h)$, using double integrals, provided one has a suitable bi-dimensional DFE to make use of.

#### 3.2 Computation of mean fitness

We assume a large number $G$ of unlinked loci, each locus having a mainland allele with a selection coeeficient drawn from the above DFE.

At some time $t$ after the onset of immigration, a locus with selection coefficient $s$ has a probability of having fixed given by eq. (1). Its contribution to the genomic fitness is thus equal to $1 + f_1(s, t)s$. The mean fitness is then the multiplicative contribution over all loci, which is computed as:

$$\bar{w}(t) = \exp \left( G \int_{-\infty}^{\infty} p(s) \ln \left(1 + f_1(s, t)s\right) \mathrm{d}s \right) \tag{8}$$

The number of loci $G$ here acts just as a scaling parameter that governs the total fitness impacts, but otherwise does not effect comparisons. Obviously, the total fitness impact can be made arbitrarily large by increasing the total number of loci. Therefore, for simplicity we presented in fig. 5 values rescaled over the total fitness impact, i.e. $G = 1$, as though the total fitness impact was kept constant and redistributed over all loci, regardless of their number. This amounts to rescaling the fitness effect of each locus as the total number of loci is made larger.

#### 3.3 Computation of migration loads, allele fixation rates and conditional FDEs

Intially, the island population is by convention at $\bar{w} = 1$, and the genetic load at time $t$ after the onset of migration is

$$L(t) = 1 - \bar{w}(t) \tag{9}$$

The pulsedness load is simply the difference between the load observed for some $n$ and the one that would be observed with continuous migration ($n = 1$), i.e. the change in genetic load caused by migration pulsedness.

At any time, the instantaneous rate of allele fixation (panel b in fig. 5) is given by:

$$\int_{-\infty}^{\infty} p(s)\left(1-f_1(s,t)\right)\frac{m}{n}u\left(\frac{n}{N+n},s\right)\mathrm{d}s \tag{10}$$

Finally, the conditional DFE, i.e. the DFE of alleles *that are currently going to fixation* (inserts in fig. 5) is obtained as the probability that an allele currently going to fixation has selection coefficient $s$, which we compute from standard probability theory as:

$$\frac{p(s)\left(1-f_1(s,t)\right)\frac{m}{n}u\left(\frac{n}{N+n},s\right)}{\int_{-\infty}^{\infty} p(x)\left(1-f_1(x,t)\right)\frac{m}{n}u\left(\frac{n}{N+n},x\right)\mathrm{d}x} \tag{11}$$

#### 4 Supplementary Figure 1: stochastic simulations with dominance

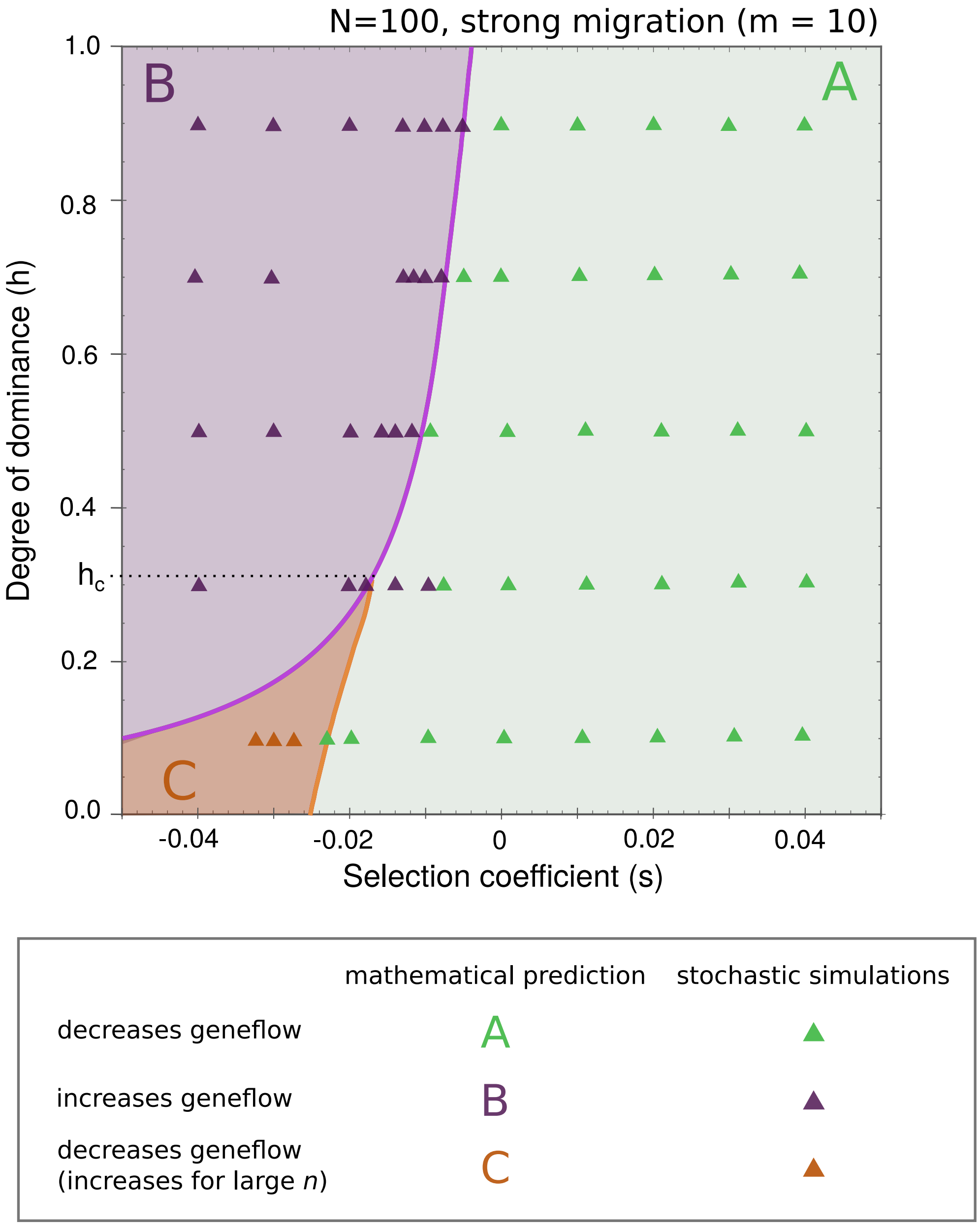

Results of stochastic simulations: same as Figure 2 in the main text, but with simulations run under the strong migration scenario ($m = 10$). As explained in the main text, the effect of pulsedness at low values of $n$ almost perfectly follows the mathematical predictions, in particular the value of $s_1$ has very good predictive power even with strong migration.
